# Trial by phylogenetics - Evaluating the Multi-Species Coalescent for phylogenetic inference on taxa with high levels of paralogy (Gonyaulacales, Dinophyceae)

**DOI:** 10.1101/683383

**Authors:** Anna Liza Kretzschmar, Arjun Verma, Shauna Murray, Tim Kahlke, Mathieu Fourment, Aaron E. Darling

**Affiliations:** The ithree institute, University of Technology Sydney, Australia; c3, University of Technology Sydney, Australia

## Abstract

From publicly available next-gen sequencing datasets of non-model organisms, such as marine protists, arise opportunities to explore their evolutionary relationships. In this study we explored the effects that dataset and model selection have on the phylogenetic inference of the Gonyaulacales, single celled marine algae of the phylum Dinoflagellata with genomes that show extensive paralogy. We developed a method for identifying and extracting single copy genes from RNA-seq libraries and compared phylogenies inferred from these single copy genes with those inferred from commonly used genetic markers and phylogenetic methods. Comparison of two datasets and three different phylogenetic models showed that exclusive use of ribosomal DNA sequences, maximum likelihood and gene concatenation showed very different results to that obtained with the multi-species coalescent. The multi-species coalescent has recently been recognized as being robust to the inclusion of paralogs, including hidden paralogs present in single copy gene sets (pseudoorthologs). Comparisons of model fit strongly favored the multi-species coalescent for these data, over a concatenated alignment (single tree) model. Our findings suggest that the multi-species coalescent (inferred either via Maximum Likelihood or Bayesian Inference) should be considered for future phylogenetic studies of organisms where accurate selection of orthologs is difficult.

## INTRODUCTION

Historically, the availability of genetic data has been the limiting factor in phylogenetic inference of evolutionary relationships. Now, the breadth of publicly available data sets generated by high throughput sequencing techniques allows for an increasingly detailed investigation into the evolutionary relationships between organisms. The quest to untangle an organism’s phylogeny is often challenging but can inform a broad range of further studies, for example epidemiology, toxicology and ecological interactions, e.g. (McTavish et al., 2017; Lewis et al., 2008; Mutreja et al., 2011; Cavender-Bares et al., 2009; Sites Jr et al., 2011).

Factors impacting phylogenetic studies range from the computational methods and availability of compute infrastructure, the methods and models applied to the data as well as the accuracy of the initial genetic data set itself. Furthermore, the practitioners themselves need to have a solid understanding of the methods, including their shortcomings.

An example of the breadth of publicly available data is the Marine Microbial Eukaryote Transcriptome Sequencing Project (MMETSP), which provides transcriptome sequences of over 650 marine eukaryotic microbes (Keeling et al., 2014). The MMETSP project focuses on a group of understudied organisms which are abundant and play vital roles in the marine environment, from geochemical cycling, to predation, to symbiosis (Gómez, 2005, 2012). This data set offers an excellent opportunity to explore the evolutionary relationships between these taxa through phylogenetics.

Central to phylogenetic inference is the existence of characters (such as nucleotides) derived from a common ancestor, which is called homology (Fitch, 2000). There are several types of homology, each differing in how the characters diverged, and determining the mechanisms through which characters have evolved is essential for choosing the correct inference model. Orthology refers to the case where the divergence of two gene copies has followed a speciation event (Fitch, 1970). Paralogs are two gene copies whose divergence is initiated by gene duplication (Fitch, 1970). Xenologs are genes which, having previously diverged from a common ancestor, have since undergone transfer between organisms through a horizontal gene transfer mechanism (Darby et al., 2016). The distinction between these cases is usually considered essential in identifying gene candidates that are informative for species evolution inference, as the selection of orthologs ensures the inclusion of a signal that is based on the speciation of the taxa examined, while selection of paralogs confounds that signal by including information that does not pertain to the speciation of the taxa (Du et al., 2019). As gene duplication and subsequent loss commonly occur over the course of evolution and speciation, the identification of genes that have orthologous relationships is more difficult than may seem apparent from the definition (Gabaldón, 2008). Importantly, identifying single copy genes does not ensure the selection of orthologous gene copies, as the gene candidates could well be from paralogous lineages where a secondary copy has been lost between the taxa. The case where paralogous homology of genes is masked by gene loss, is termed pseudoorthology (Koonin, 2005).

Once candidate genes have been identified, there are further issues that can arise and impact the veracity of the phylogenetic inference. Two common, well characterized types of errors are random (sampling error) and systematic errors. The former arises from the data, as individual gene histories may differ to the species tree. With a small number of genes, this error can reduce the confidence (through node support values) of the topology, and in extreme cases can entirely skew the inference away from resolving a good approximation of a species tree. Increasing the number of genes directly reduces the impact this error has on the analysis (Philippe et al., 2004; Heath et al., 2008).

Conversely, systematic errors arise due to the misspecification of the model used for the inference, leading to an incorrect species tree topology. In this case, an increase in data set size can exacerbate systematic errors rather than reduce them as would happen with random errors (see Box 1). In the presence of this type of systematic error, the resulting inference can be positively misleading, with high clade support values for the incorrect tree topology, obfuscating the presence of the error (Jeffroy et al., 2006; Roch and Steel, 2015; Kubatko and Degnan, 2007).

**Box 1: Statistical nomenclature & errors this study seeks to address**

For in-depth explanations see (Yang, 2014).

- **Potential statistical error types:**
  1. random. Sampling-based error which decreases and approaches zero as the size of the data set approaches infinity.
  2. systematic. Arises from incorrect model assumptions or problems with the model itself. Error type persists and increases as data set size approaches infinity. If strong, can override true phylogenetic signal.
- **Incomplete lineage sorting (ILS):** discordance of gene evolutionary history with the species evolutionary history causing the phylogenetic species tree to be incorrectly inferred. Difference in the topology of a gene tree compared to the species evolution can arise from the divergence of those orthologs prior to the species divergence, where in effect the ancestral populations contain two or more already diverged copies of the gene across one or more species divergence points. Another mechanism is the introduction of a copy of the gene which is not based on ancestral inheritance (xenology), such as horizontal gene transfer or hybridization.
- **Long branch attraction (LBA):** placement of two heavily divergent but non-monophyletic sequences with each other. The model is unable to extract the correct evolutionary signal due to the number of mutations that have occurred, so places the two taxa together. Also called the Felsenstein zone.

In summary, common problems in carrying out a species tree inference arise from:

1. Selection of paralogs (including pseudoorthologs). If genes with different evolutionary histories are selected, and if this violates the phylogenetic model, the inferred tree may not accurately reflect the history of any of the individual genes or that of the species;
2. Concatenation of genes. Can be a statistically inconsistent estimator of the species tree due to incomplete lineage sorting (ILS) and concatenation acts as an imperfect estimator of species tree topology (Roch and Steel, 2015);
3. Inference of model adequacy from bootstrap values. Kubatko and Degnan (2007) demonstrated high bootstrap support under maximum likelihood (ML) inference for incorrect species trees with concatenated gene sets as input (Kubatko and Degnan, 2007). As high bootstrap values are often used as an indicator for robust species topology resolution, this fallacy is particularly problematic if the reader/operator is unfamiliar with the statistical phenomenon.

In this study, we explored the application of data analysis techniques which attempted to mitigate several of the pitfalls in species tree inference, beyond what has previously been applied in the study of protist phylogenetics. The sequence data was prepared using a workflow that assembled RNA-seq data sets, identified and extracted single copy genes across input taxa, and aligned selected genes ready for Bayesian inference (BI) phylogenetics. Next, we evaluated the impact of model and data selection on the resulting phylogenetic inference. Finally, we applied the methodology to a group of organisms notorious for their extensive paralogy - the Gonyaulacales (phylum: Dinoflagellata) (see box 2 for further information on the dinoflagellates). We present a phylogenetic inference of the Gonyaulacales generated under the multi-species coalescent (MSC) and compare the topology to inferences with commonly used methodologies.

**Box 2: Who/what are the Gonyaulacales?**

The Gonyaulacales are an order within the super-phylum Alveolata and sub-phylum Dinoflagellata, which are an ancient eukaryotic lineage (Moldowan and Talyzina, 1998). They play a role in several important ecological processes in aquatic environments where they cover a diverse array of niches such as symbionts, parasites and autotrophs. Some taxa can cause harmful algal blooms through proliferation (by restricting light and nutrient availability to other organisms) and/or neurotoxin production (e.g. causing paralytic shellfish poisoning, ciguatera fish poisoning) (Murray et al., 2016). Dinoflagellates possess large genomes (estimated size range 1.5 to 185 Gbp), with extensive paralogy and repetitive short sequences (Casabianca et al., 2017). In particular paralogy has proven problematic for efforts investigating the genetic content and structure of the dinoflagellates, as this feature has prevented the assembly of genomes apart from draft genomes from symbiodiniacean taxa which posses some of the smaller genomes (Shoguchi et al., 2013; Lin et al., 2015; LaJeunesse et al., 2018). Gene duplication, loss, and cDNA recycling is rife within these organisms, therefore they have likely undergone complementary gene deletion events (Slamovits and Keeling, 2008; Murray et al., 2015; Shoguchi et al., 2018). For a review on the genetic features of dinoflagellates see Murray et al. (2016). While the evolutionary relationship of most orders within the dinoflagellates has been inferred with consistently high support values, one order has often escaped elucidation - the Gonyaulacales. As neurotoxin production, which can accumulate up the food chain, is prevalent in this order, the evolution of the order is of interest to provide a frame of reference for future investigations into how the toxins have evolved (Shalchian-Tabrizi et al., 2006; Zhang et al., 2007; Saldarriaga et al., 2004; Hoppenrath and Leander, 2010; Murray et al., 2005).

## METHODS

### Culture conditions

Cultures were isolated from locations as per Table S1 and clonal cultures established by micropipetting single cells through sterile seawater as described in in (Kretzschmar et al., 2017). Clonal cultures were maintained in 5x diluted F/2 medium (Holmes et al., 1991) and maintained at temperatures indicated in Table S1.

### RNA isolation, library preparation and sequencing

*Gambierdiscus* spp. and *Thecadinium kofoidii* were harvested during late exponential growth phase by filtration onto 5 *µ*m SMWP Millipore membrane filters (Merck, DE) and washed off with sterile seawater. Cells were pelleted via centrifugation for 10 minutes at 350 rcf. The supernatant was decanted and 2ml of TRI Reagent (Sigma-Aldrich, subsidiary of Merck, DE) was added to the pellet and vortexed till resuspended. Samples were split in two and transferred to 1.5ml eppendorf tubes. Cellular thecae were ruptured by three rounds of freeze-thaw, with tubes transferred between liquid Nitrogen and 95 *°*C. RNA was extracted as per the protocol for TRI Reagent (Rio et al., 2010). RNA eluate was purified with the RNeasy RNA clean up kit RNeasy Mini Kit (Qiagen, NL) as per protocol. DNA was digested with TurboDNAse (Life Technologies, subsidiary of Thermo Fischer scientific, AU). RNA was quantified with a Nanodrop 2000 (Thermo Scientific, Australia) and frozen at −80 *°*C until sequencing. The quality of samples was assessed via an Agilent 2100 Bioanalyzer at the Ramaciotti Center (UNSW, AU) and the libraries were prepared using TruSeq RNA Sample prep kit v2 (Illumina, USA). Paired-end sequencing was performed with a NextSeq 500 High Output run at the Ramaciotti Center (UNSW, AU) with 75bp read length for *G. holmesii* and *G. lapillus*; and 150bp read length for *G. carpenterii, G. polynesiensis* and *T.kofoidii*.

#### Publicly available transcriptome libraries

From NCBI, the *Gambierdiscus excentricus* VGO790 transcriptome was downloaded via the accession ID SRR3348983 (Kohli et al., 2017), while *Coolia malayensis, Ostreopsis ovata, Ostreopsis rhodesae* and *Ostreopsis siamensis* transcriptomes were downloaded via the accession IDs SRR9044102, SRR9046040, SRR9047231 and SRR9038703 respectively (Verma et al., 2019). Accession numbers are provided in table 1. RNA-seq libraries for all remaining transcriptomes were generated by, and downloaded from, the MMETSP (Keeling et al., 2014).

**Table 1.**
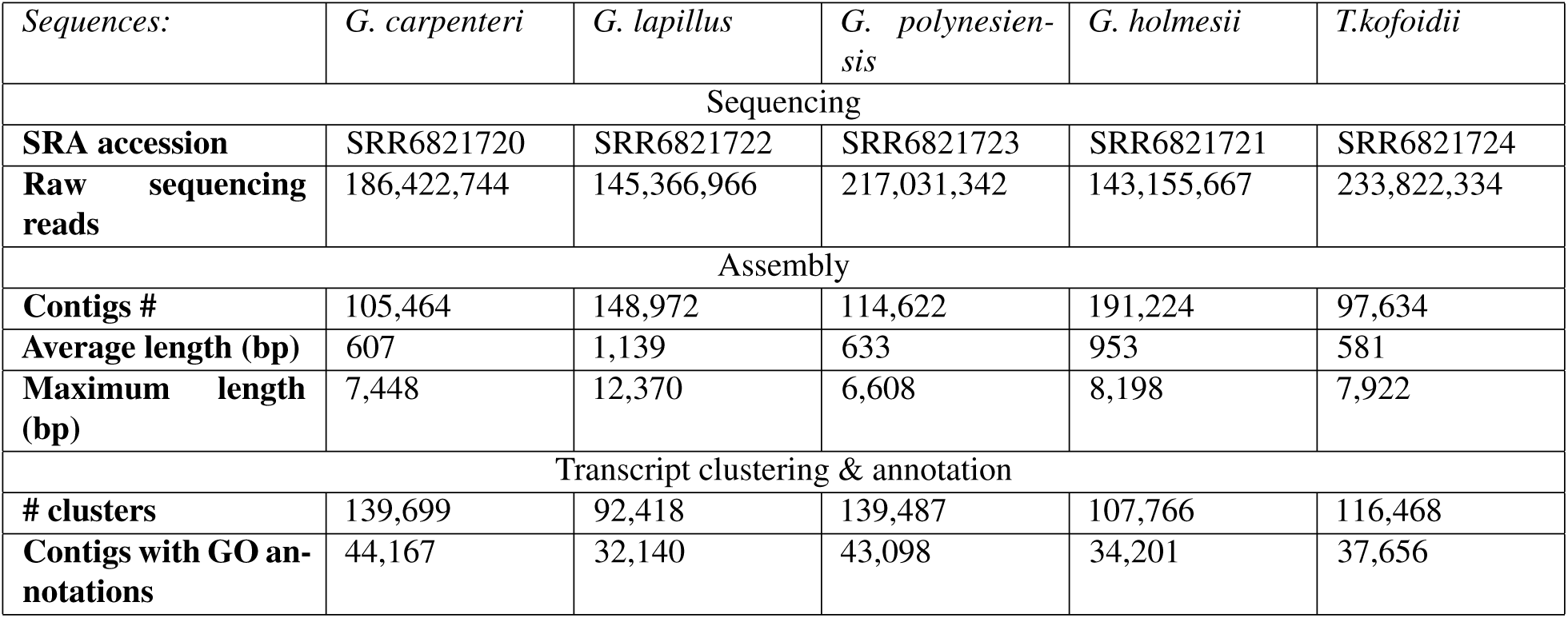
Summary of transcriptome sequencing and assembly statistics.

### Transcriptome processing scripts

The workflow was separated into two parts. See section Implementation for script details.

#### Transcriptome assembly

Individual RNA sequencing libraries were processed through FastQC (Andrews, 2010) for quality metrics, sequences were trimmed with Trimmomatic (`LEADING:3 TRAILING:3 SLIDINGWINDOW:4:5 MINLEN:25`) (Bolger et al., 2014) and assembled with Trinity v2.4.0 (default settings for paired end libraries) (Haas et al., 2013). Assemblies were then processed with BUSCOv2 with the protist specific library (Simão et al., 2015). The RNA libraries with 150bp reads generated as part of this study were also subjected to Digital Normalization (Brown et al., 2013) prior to assembly, to reduce data set size by removing highly similar sequences, which were then used for downstream analysis.

#### Construction of multiple sequence alignments

The BUSCOv2 output from all transcriptomes from the previous step formed the input for identification of single copy genes and construction of multiple alignments. Any genes that BUSCOv2 identified as single copy and were present in at least 75% of the transcriptomes were indexed, the corresponding contig extracted from the assemblies, aligned with hmmer3.1b2 (Eddy and Wheeler, 2015) and unaligned regions trimmed. If several candidate sequences were processed for the same organism, a warning message in the terminal window alerted the user before proceeding. The output for this section was used as a basis for single copy gene phylogenetic inferences in subsequent sections.

### Assembly analysis

Contigs from assemblies were clustered with CD-HIT with the flags `-T 10 -M 5000 -G 0 –c 1.00 -aS 1.00 -aL 0.005` (Fu et al., 2012). Protein coding regions within the clusters were predicted with Transdecoder (Haas and Papanicolaou, 2016). Amino acid clusters were clustered again with CD-HIT with the flags as previously except `-c 0.98`. Protein sequences were analyzed with interproscan v5.27 with local lookup server (Quevillon et al., 2005).

### Phylogenetic inferences

#### Ribosomal DNA based inference

Ribosomal DNA (rDNA) sequences for the small subunit (SSU) region as well as the D1-D3 large subunit (LSU) region were acquired from NCBI (Coordinators, 2017) and the SILVA rRNA database project (Quast et al., 2013), accession IDs in Table S3. Individual genes were aligned using MUSCLE (Edgar, 2004) for a maximum of 8 iterations and then were concatenated in Geneious v11.3 (Kearse et al., 2012). ML phylogenies were inferred using RaxML (Stamatakis, 2014) with the model GTRGAMMA and with 100 bootstrap replicates.

#### Inference of concatenated single copy genes

Amino acid substitution model selection was carried out with ProtTest3 with the Bayesian Information Criterion as well as the log likelihood (Darriba et al., 2011; Guindon and Gascuel, 2003). The best-fit model for the data set identified by both criteria was VT followed by LG, however neither are available in BEAST2 so the third best model, WAG, was chosen for analysis.

##### Maximum likelihood with concatenated sequences

ML inference was run as described in the previous section, with the PROT, GAMMA and WAG flags.

##### Bayesian inference with concatenated sequences

BI was run in BEAST2 with the Gamma site model with 4 discrete categories under the WAG substitution model (Whelan and Goldman, 2001). A local random clock was used under the birth-death model 3,000,000 million chains.

##### Bayesian probability under the MSC

BI of the species tree was carried out under the *BEAST2 model in BEAST2 (Bouckaert et al., 2019). The analysis was performed with the WAG amino acid substitution model (Whelan and Goldman, 2001) and with a Gamma distribution with four rate categories. A random local clock was employed (Drummond and Suchard, 2010). Posterior distributions of parameters were approximated after 300,000,000 generations of MCMC, subsampled every 5,000 generations with a burn-in of 15%. The inference was run four times to evaluate convergence of parameters, then log and tree files (without burn-in) were merged.

### Marginal likelihood analysis

We estimated the marginal likelihood of the data under the coalescent (i.e. concatenated alignment) and the MSC (*BEAST) models to compare their fit. We used the stepping stone algorithm by Xie et al. (2011) along a path of 30 power posteriors. The *β* values are set equal to the quantiles of the beta distribution with shape parameter *α* = 0.3 and *β* = 1, as recommended by Xie et al. (2011).

#### Generation of figures

Tanglegrams were generated with Dendroscope v3.5.9 (Huson et al., 2007); images were edited in GIMP (Gimp, 2008) to improve readability.

#### Implementation

The analysis workflow in section “Transcriptome processing scripts” was constructed as a Nextflow work-flow (Di Tommaso et al., 2017) and is available on Github at https://github.com/hydrahamster/gonya phylo. Packages within the scripts are written in bash, Python 2.7 (Stevens and Boucher, 2018) and pandas (McKinney, 2010). Source code for the scripts is provided under an open source license. The scripts (1) assemble RNA-seq data sets, (2) identify and extract single copy genes across input taxa with extensive paralogy, and (3) align selected genes in preparation for phylogenetic analysis. The data sets were processed on a Genomics Virtual Lab (GVL) (Afgan et al., 2015) instance in the NeCTAR cloud. Phylogenetic analyses were carried out on the University of Technology Sydney’s High-performance computing cluster (HPCC) and were accelerated using BEAGLE (Ayres et al., 2011) on the GPU. GPU processing units were either Nvidia Tesla K80 or a Tesla P100.

## RESULTS

Assemblies, annotation files, BUSCOv2 output, single-copy gene alignments and single copy gene MSC BI trace files generated in this study are available on Zenodo doi: 10.5281/zenodo.2576201

### Transcriptomes overview

RNA-seq libraries generated in this study are available in the NCBI sequence read archive (SRA) under the project ID SRP134273. Sequencing of transcriptomes for *Gambierdiscus* spp. and *T.kofoidii* generated data sets ranging in size from 143,155,667 to 233,822,334 reads, resulting in 97,634 to 191,224 assembled contigs (table 1). Clusters with gene ontology (GO) annotations made up 30.9% to 34.8% of the total clusters.

### Single copy gene search with BUSCOv2

Assemblies were searched with BUSCOv2 for 234 candidate single copy genes and homologs to these single copy genes were extracted. The single copy genes acquired through the BUSCO HMMER libraries curated for protists are reported in Table S2, as well as accession numbers and identifiers for each transcriptome. The alignments are available on Zenodo, with the BUSCO gene IDs included in the alignment name.

### Phylogenetic inference

Support for branches was interpreted as follows, for ML and BI, respectively: 100%/1.0 was considered fully supported, above 90%/0.9 was very well supported, 80%/0.8 and above was interpreted as relatively well supported and above 50%/0.5 was considered weakly supported. Below 50%/0.5 was considered unsupported. As *Azadinium spinosum, Dinophysis acuminata* and *Karenia brevis* are members of different orders (Dinophyceae ordo incertae sedis, Dinophysiales & Gymnodiniales respectively) and are consistently placed outside of the Gonyaulacales in phylogenetic analyses, their placement as an outgroup was considered a given for this study. Therefore, the branch separating these taxa from others was used to root ML trees in subsequent analyses where rooting was required for tree layout in visual comparisons.

#### rDNA based phylogeny

All nodes were supported, with a range of certainty (Fig. 1). Species within the genera *Gambierdiscus* and *Ostreopsis* resolved with their sister species with full support. Within the *Gambierdiscus* clade, nodes were either weakly supported or fully supported. The two species of *Alexandrium* resolved as well supported closest relatives, but did not form an individual clade. Deeper nodes were supported but with less certainty than the nodes near the tips. Two distinct clades were be observed from the topology: One including *Alexandrium, Coolia* and *Ostreopsis*; another with only *Gambierdiscus*. Sister to these clades, in descending order, was *Pyrodinium, Ceratium* and *Gonaulax, Protoceratium* and *Thecadinium*. The outgroup were relatively well supported and included *Crypthecodinium*. Support for deeper nodes varied from weak to well supported.

**Figure 1.**
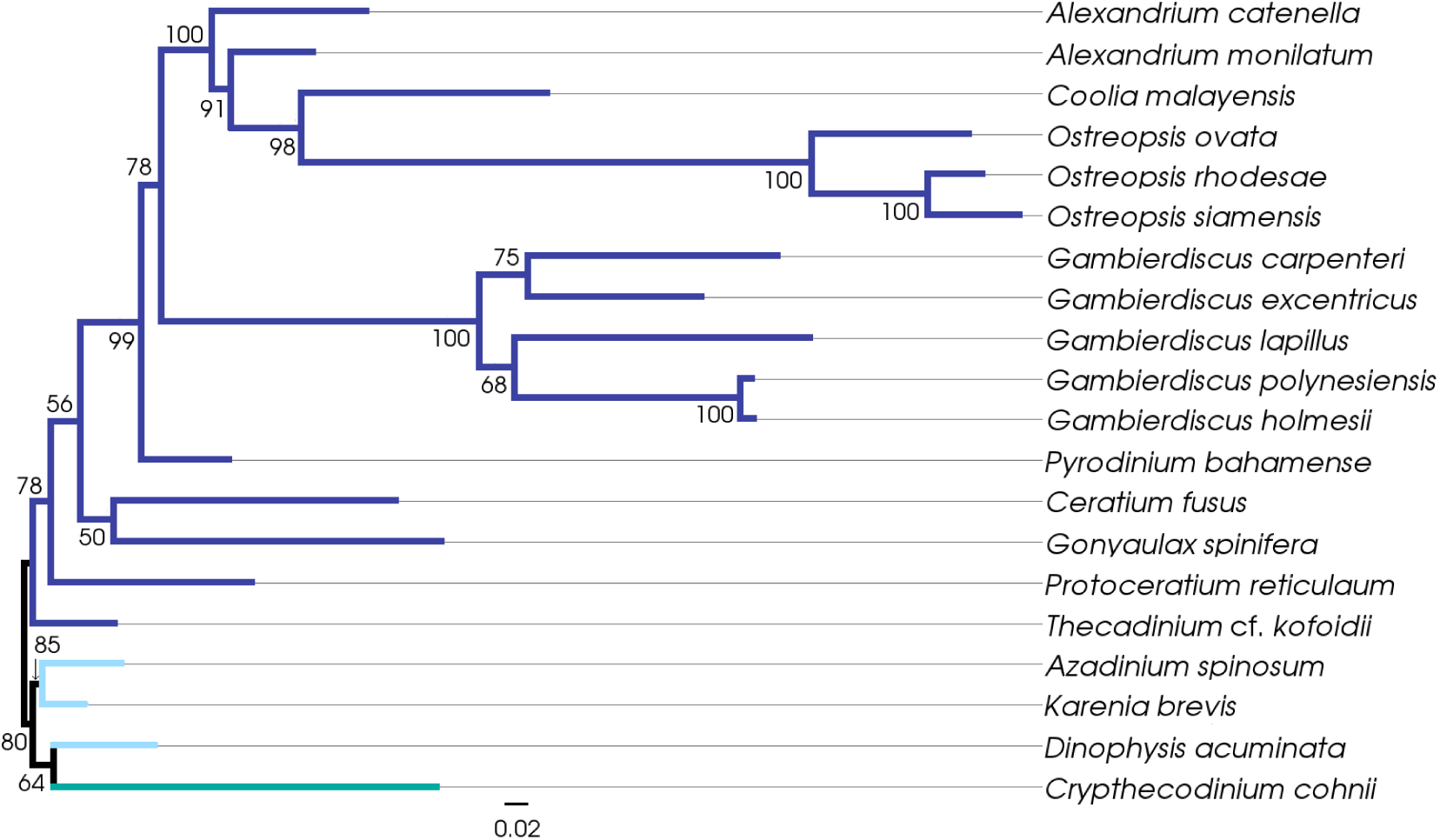
Maximum likelihood phylogenetic inference of ribosomal DNA genes. Concatenation of small subunit rDNA and D1-D3 region large subunit rDNA. Accession numbers for concatenated genes in Table S3. Gonyaulacales (n=16) in purple, outgroups (n=3) in light blue and taxa *incertae sedis* (n=1) in teal. The topology was rerooted on the branch separating outgroup taxa with the Gonyaulacales. The scale represents the expected number of substitutions per site.

#### Concatenated single copy gene based phylogeny inferred with ML

All nodes except one within the *Gambierdiscus* species cluster were relatively well supported (Fig. 2). Species of the genera *Alexandrium, Gambierdiscus* and *Ostreopsis* clustered as individual clades with their sister species. The topology showed three distinct, well supported clades: One encompassing *Alexandrium, Coolia* and *Ostreopsis*; another which only contained *Gambierdiscus*; and one which includes *Pyrodinium, Gonyaulax* and *Protoceratium*. Sister to these clades is *Thecadinium*, followed by *Ceratium*. The split of the outgroup was fully supported, while the internal nodes were very well supported. *Crypthecodinium* was placed within the outgroup, sister to *Karenia*. Other deeper nodes were well supported.

**Figure 2.**
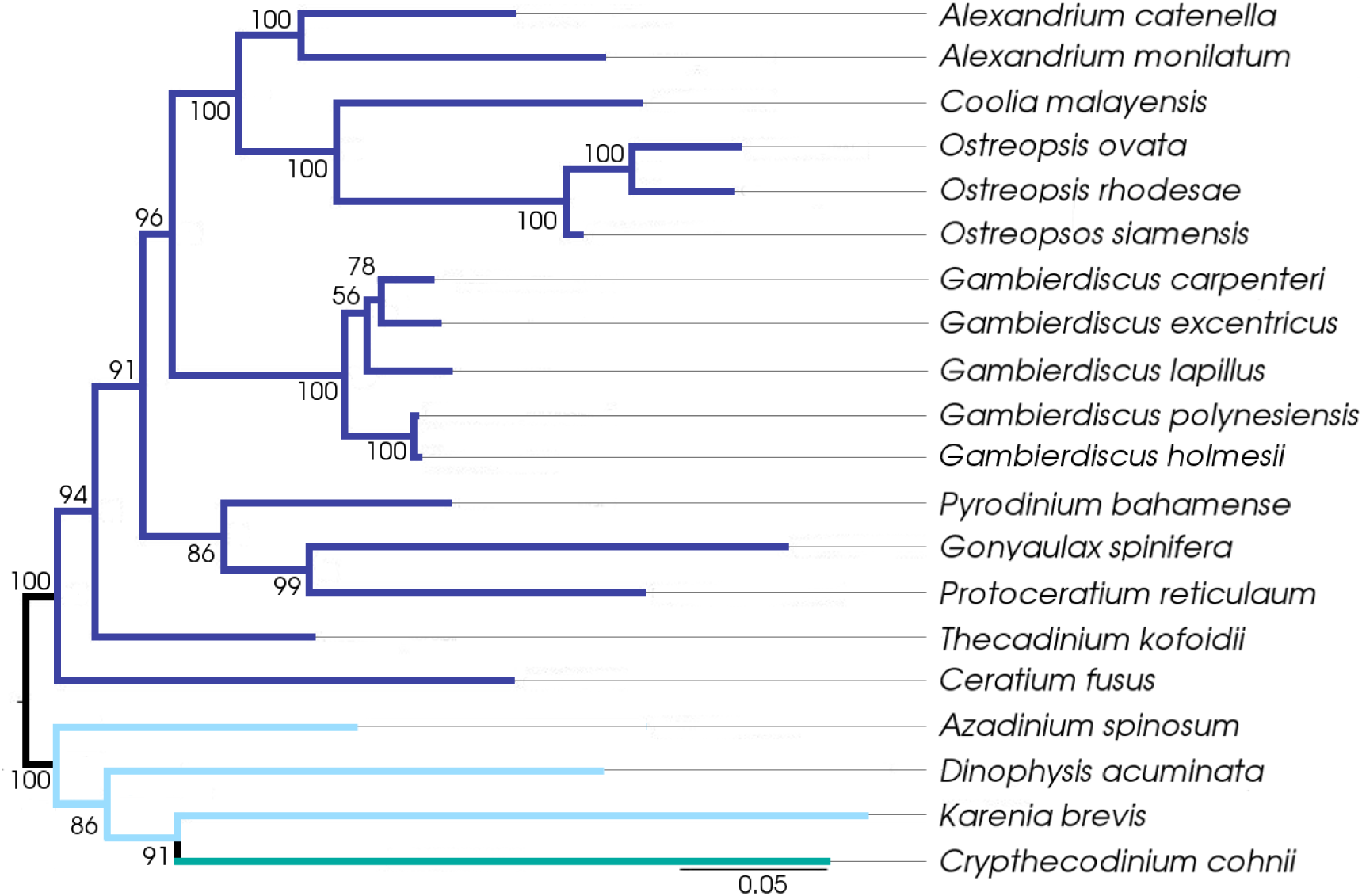
Maximum likelihood phylogenetic inference of concatenated single copy gene set (62 single copy genes from 20 taxa). Gonyaulacales (#16) in purple, outgroups (#3) in light blue and taxa *incertae sedis* (#1) in teal. Topology was rerooted on the branch separating the outgroup taxa from the Gonyaulacales. The scale represents the expected number of substitutions per site.

#### Concatenated single copy gene based phylogeny inferred with BI

All nodes resolved with full support, except one node within the genus *Gambierdiscus* which was very well supported as well as an internal node within the outgroup clade (Fig. 3). The species in the genera *Alexandrium, Gambierdiscus* and *Ostreopsis* were monophyletic with full support. The overall topology of the Gonyaulacales was resolved as three clades with *Thecadinium* and then *Ceratium* as ancestral lineages. *Alexandrium, Coolia* and *Ostreopsis* clustered together, followed by *Gambierdiscus* on their own in a sister clade. The third clade encompassed *Gonyaulax, Protoceratium* and *Pyrodinium*.

**Figure 3.**
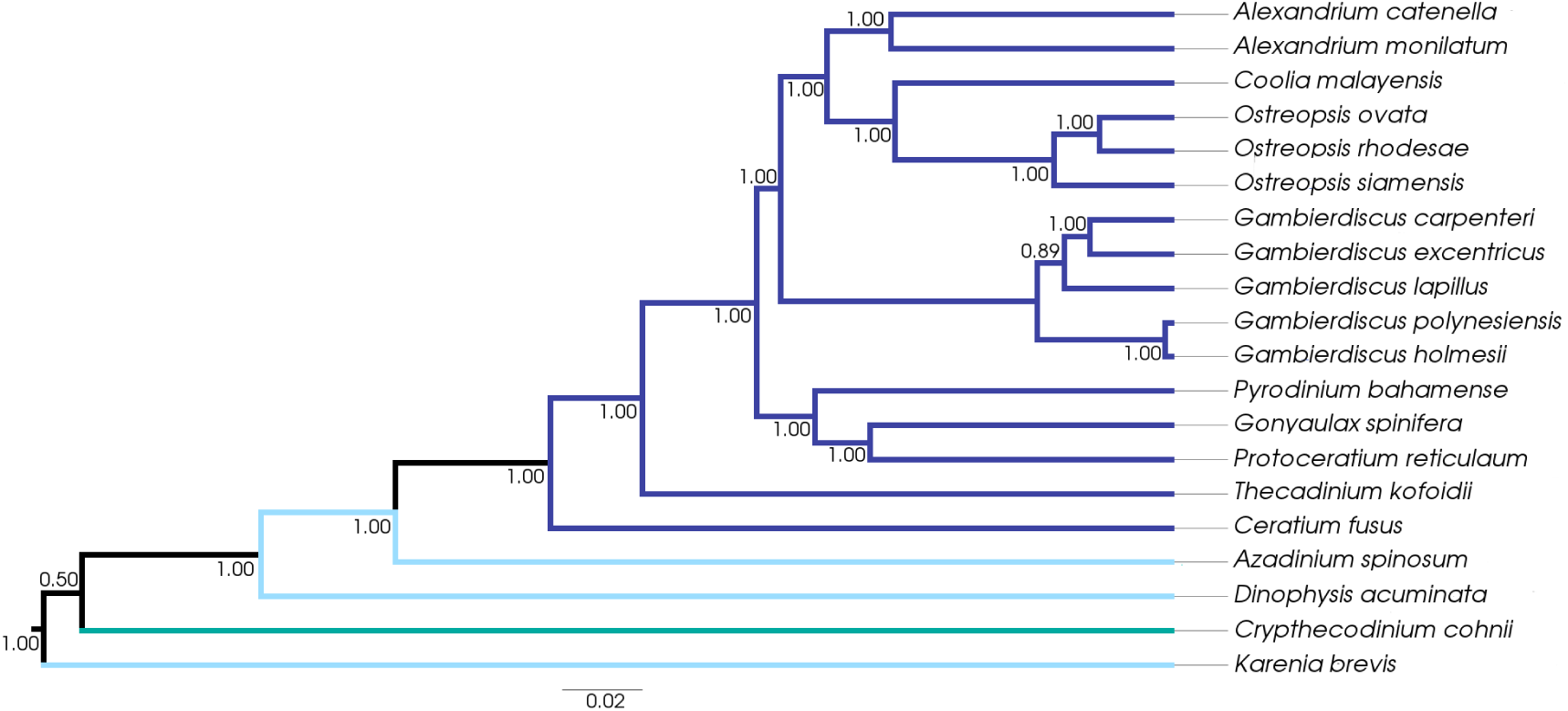
Bayesian phylogenetic inference of concatenated single copy gene set (62 single copy genes from 20 taxa). Gonyaulacales (#16) in purple, outgroups (#3) in light blue and taxa *incertae sedis* (#1) in teal. The scale represents the expected number of substitutions per site.

#### Single copy gene based phylogeny under MSC

Species of *Alexandrium, Ostreopsis* and *Gambierdiscus* were either well or fully supported within their genus clades (Fig. 4). The topology within the Gonyaulacales resolved into three clades: one fully supported encompassing *Alexandrium, Coolia* and *Ostreopsis*; a well supported clade with *Gambierdiscus* and *Pyrodinium*; and a weakly supported clade including *Ceratium, Gonyaulax, Protoceratium* and *Thecadinium*. The outgroup taxa clustered together with high support. *Crypthecodinium* was placed as a sister taxon to the outgroup. Other deeper nodes were well or fully supported.

**Figure 4.**
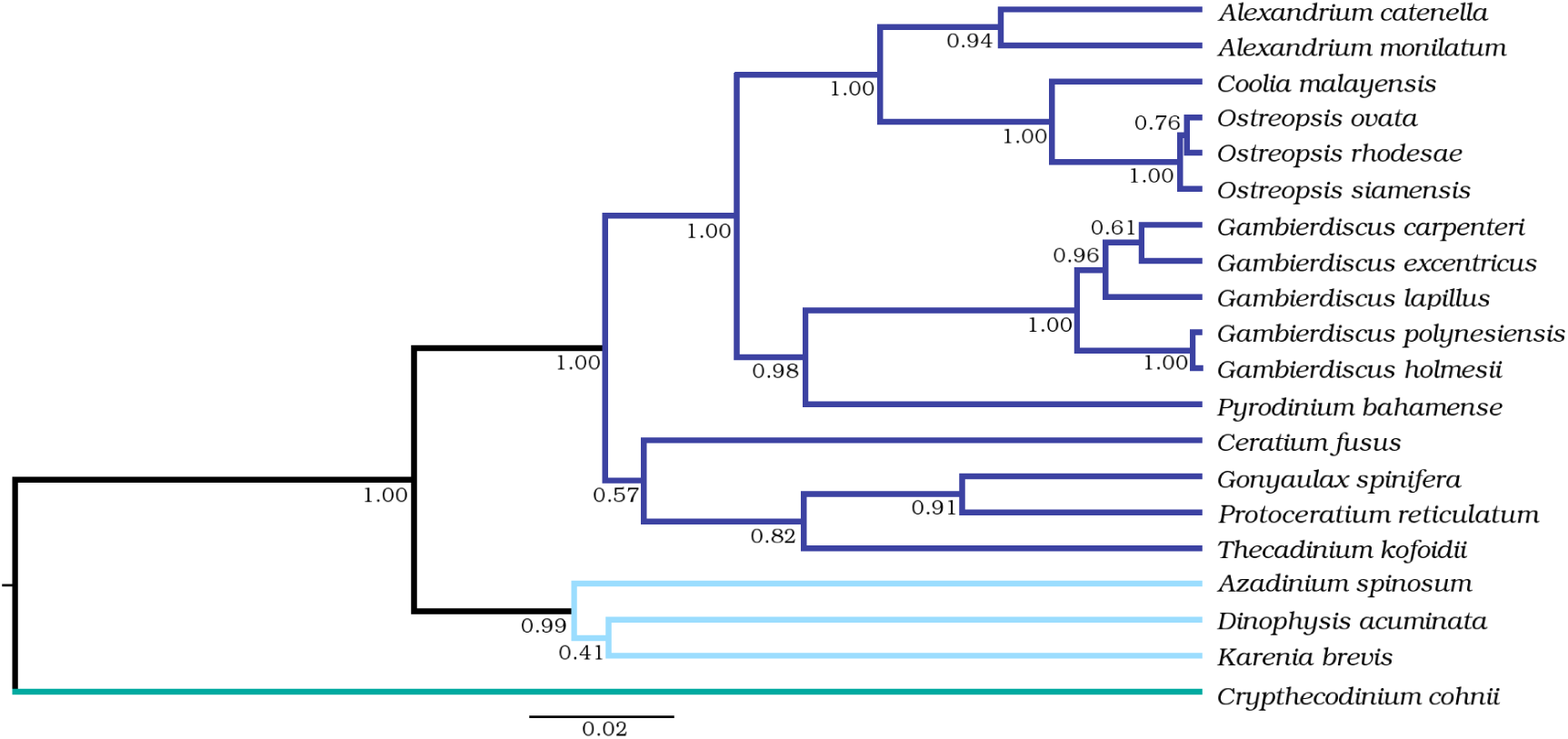
Bayesian phylogenetic inference of a Gonyaulacales species tree under the MSC model with 62 single copy genes from 20 taxa. Gonyaulacales (#16) in purple, outgroups (#3) in light blue and taxa *incertae sedis* (#1) in teal. The scale represents the expected number of substitutions per site.

## DISCUSSION

Phylogenetic inference is a fundamental approach for exploration of evolutionary relationships between organisms, with applications in pathology, ecology, investigating adaptive traits and many more (Heath et al., 2008). Advances in sequencing technologies have seen an increase in high throughput sequencing initiatives such as MMETSP, which revealed the genomic diversity of a relatively uncharacterized group of marine microbial eukaryotes (Keeling et al., 2014). However, the methodologies used for investigating the evolutionary relationships using this type of genome-scale data remain an obstacle, as the choice of input data and method employed influences the outcome of the inference. In particular, the effects of paralogs and pseudoorthologs (hidden paralogy) are particularly problematic as they can lead to incorrect inference with classic phylogenetic methods. To address this, a synopsis on a method for single copy gene extraction, and synthesis of phylogenetic inference model availability and selection is presented in this study - as well as possible shortcomings of the parameters and methods selected.

Dinoflagellates are notorious for their large genomes with suspected whole or partial genome duplication and potential cDNA retro-insertion into the genome (Van Dolah et al., 2009; Beauchemin et al., 2012; Slamovits and Keeling, 2008; Hou and Lin, 2009; Lin, 2011). This can lead to unusually high gene copy numbers and extensive paralogy. With this in mind, the Gonyaulacales (an order within the dinoflagellates, see box 2) represented a good case study for examining the impacts of paralogy on phylogenetic inference. This study presents the first species tree for the Gonyaulacales that has been inferred with a method robust to paralogy, including hidden paralogy.

The phylogenetic inference for Gonyaulacales that resulted from the workflow we developed, which incorporates several of the most recent innovations in analytical methodology, resolved within-genus relationships well and showed high posterior probability support throughout the species tree (Fig. 4). The inferred species tree topology followed a broad revised taxonomic classification of the Gonyaulacales based on morphological characteristics (Hoppenrath, 2017) and was used as a point of comparison to results from other commonly employed methods in later sections. The scripts which form the basis of this study are publicly available through github and the single copy gene alignments used to infer the species trees, as well as the XML input and log files for the *BEAST2 runs, are available on zenodo (doi: 10.5281/zenodo.2576201). Our study was designed to be transparent and reproducible for those with basic programming skills.

### Considerations for data set selection and pre-processing

#### Quantity of taxa in phylogenetic inference

Two phenomena that can confound the veracity of conclusions drawn from phylogenetic inference are ILS and LBA. The impact of ILS on phylogenetic inference has been explored through simulated data sets with a known species tree. When species have recently diverged, sampling more individuals per species can improve resolution of the species tree. However, when the species divergences are older, as is the case here, using more gene loci per species yields greater resolving power than sampling more individuals per species (Maddison and Knowles, 2006).

LBA can arise if some species have disproportionately high substitution rates, leading to the presence of long and short branches in the phylogenetic tree (Liu et al., 2014). The risk of LBA artefacts can be reduced by denser taxon sampling to break up long branches and ensuring that the models specified are appropriate (Heath et al., 2008). The Gonyaulacales data set in this study included a single representative species per genus, with the exception of *Alexandrium, Gambierdiscus* and *Ostreopsis*. This resulted in some genera on long branches (eg. fig. 4: *Ceratium fusus* & *Pyrodinium bahamense*) indicative of a proportionally large number of genetic changes to their closest relative. This tree shape was consistent with sparse taxon coverage and can lead to LBA artefacts (Heath et al., 2008). To investigate the presence of ILS and as a topological comparison to the BI and ML inferences, a neighbor-joining (NJ) inference was run as well (Phylip with Protdist JTT matrix and neighbor packages (Felsenstein, 2005)). The rationale for evaluating this method was that NJ can recover an accurate species topology despite ILS in cases where ML would fail (Mendes and Hahn, 2017). However NJ is more susceptible to LBA than ML or BI methods. The resulting topology was so anomalous, with out and in-groups clustering together as well as negative length branch lengths, that we chose to exclude it from further discussion. Both BI and ML are more robust to the effects of LBA than NJ, where BI tends to outperform ML especially if the latter is performed conjunction with concatenation (Kubatko and Degnan, 2007; Roch and Steel, 2015).

#### Quality of transcriptome assemblies

Publicly available data sets may have been generated with a variety of different methods, and their resulting quality can be highly variable, so an initial quality assessment step is essential. In the time since the MMETSP data sets were made available, several studies have utilized a broader range of taxa to explore evolutionary stories involving the Gonyaulacales. However, these have relied on the assemblies supplied as part of the project. The stringency for quality trimming of RNA-seq libraries prior to assembly plays a role in determining the number of unique contigs recovered and the subsequent assembly quality of transcriptomes. Regarding the transcriptome assembly method, Johnson et al. (2018) evaluated the publicly available assemblies from MMETSP using BUSCO scores, compared to processing and re-assembly with Trinity (Johnson et al., 2018). Johnson et al. (2018) demonstrated that while the raw data available from the MMETSP project is an excellent resource, the assemblies available as part of the project are of a lower quality than what can be achieved with current methods (Johnson et al., 2018). Another factor in assembly quality is RNA-seq data processing prior to assembly, especially trimming. High stringency is usually favored, however MacManes (2014) found that this can be detrimental to the assembly and the quality cut off scores used in the present study were based on those recommendations (MacManes, 2014). In short, the trimming and assembly pipeline used for the assemblies available as part of MMETSP is no longer state-of-the-art and this is reflected in the quality comparison conducted by Johnson et al. (2018). To address this problem, we developed a workflow implementation of current best-practice transcriptome assembly methods as part of this study.

##### Assembly parameters

Trinity was chosen as the assembler for this study based on the findings of Honaas et al. (2016), in which Trinity was one of the top performing assemblers for *de novo* transcriptomes as tested with *Arabidopsis thaliana*. Further, Trinity performed well for identifying isoforms of genes and excelled at assembling highly expressed genes (Honaas et al., 2016). Conversely, Cerveau and Jackson (2016) found that Trinity, CLC Bio and IDBA-Tran assemblies all contain errors introduced by the assembly algorithms. Using a combination of all three assemblers yielded a final assembly closer to biological reality than any individual assembler, when no reference genome is available (Cerveau and Jackson, 2016). As our present study used Trinity exclusively, it may be subject to the type of errors found by Cerveau and Jackson (2016) which could affect downstream analysis.

#### Selection of paralogs to infer species evolution

Inclusion of genes which diverged through a process other than speciation events, such as paralogs, violates the assumptions of most commonly used phylogenetic models which assume all genes analysed have an orthologous relationship. This study sought to mitigate the issues arising from paralogs by identifying and using single copy genes and using a phylogenetic inference method that is robust to the presence of pseudoorthologs (hidden paralogs). Single copy genes were identified via the curated BUSCO gene collection and software. As BUSCO uses lineage specific profile HMM libraries designed to target single copy genes, and the output distinguishes between single copy genes and duplications, it presents a method for reliably screening for single copy genes for phylogenomics (Waterhouse et al., 2017). Despite the known effect of paralogy on phylogenomic analyses, the first study to address this issue for species inference within the dinoflagellates by using single copy genes as input for the phylogenetic inference was only published in 2017 (Price and Bhattacharya, 2017). A second study by Stephens et al. (2018) expanded on the dataset by Price and Bhattacharya (2017) but used the same methodology for single copy gene extraction and inference, so we compares the phylogeny by Price and Bhattacharya (2017) to the one presented here as it represented a comprehensive baseline phylogenomic analysis that also includes the order Gonyaulacales.

The phylogenies inferred by Price and Bhattacharya (2017) and by our study resulted in markedly different topologies. Specifically, the placement of two sister taxa (Fig. 5) are noteworthy: Price and Bhattacharya (2017) placed *Alexandrium* spp. as the closest genus to *Gambierdiscus*, while our study placed *Pyrodinium* as the sister to *Gambierdiuscus*. Interestingly, one of the few points of difference between the Price and Bhattacharya (2017) and the Stephens et al. (2018) inference topologies was that the latter placed *Pyrodinium* as the sister genus to *Gambierdiscus* too. Similarly, in Price and Bhattacharya (2017) the *Azadinium* is part of the Gonyaulacales, while this study firmly places this genus as an outgroup with *Dinophysis* spp. and *Karenia* spp. Given that some *Gambierdiscus* spp. as well as *Azadinium* spp. produce toxins that cause severe fish and shellfish poisoning it is of high importance for the analysis of toxin evolution to infer the phylogenetic relationships of these and closely related taxa (Pawlowiez et al., 2014). Potential factors that may have impacted the present study and explain the differences between the two phylogenies are discussed in detail below. Additionally, factors that may have impacted the phylogeny published by Price and Bhattacharya (2017) which could also explain the observed differences are (i) older assembly methods used in the MMETSP data set, (ii) the concatenation of genes for the alignment as well as (iii) the use of a ML estimation method. In our opinion, especially the use of concatenated alignments in conjunction with ML inference methods (as discussed previously) makes the study published by Price and Bhattacharya (2017) susceptible to the effects of pseudoorthologs (e.g. hidden paralogs). Unfortunately, a more rigorous comparison between the two approaches was not possible as the methodology for the identification of single-copy genes was neither reported by Price and Bhattacharya (2017) nor available on request.

**Figure 5.**
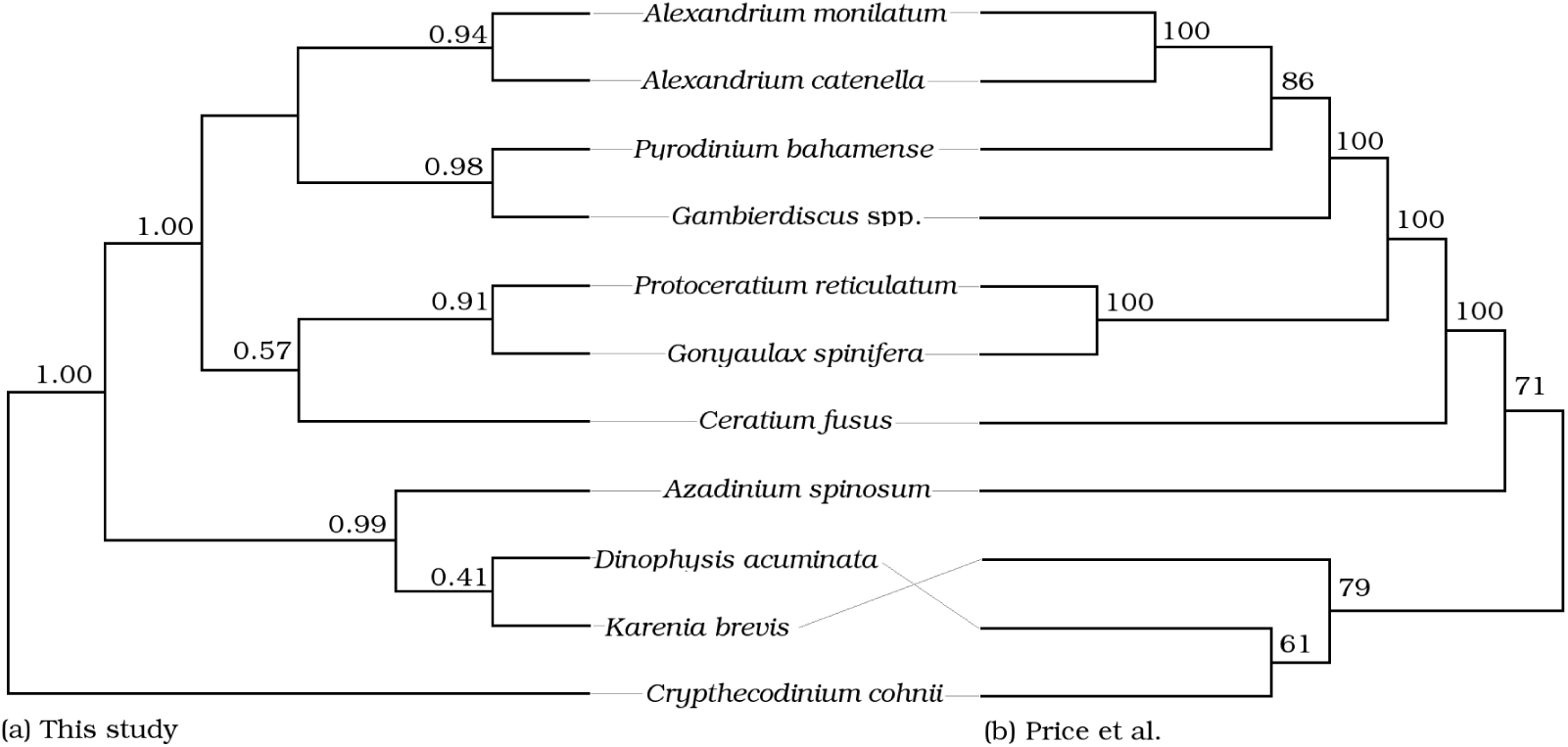
Tanglegram of the single copy gene topologies presented in (a) this study under MSC; and (b) concatenated by Price and Bhattacharya (2017). Taxa not common to either study are not shown due to the reduced topologies from the original studies, closest PP or BS to branch split were included.

#### Model selection for inference

The issue of model choice is an important one, as the choice of model can heavily influence the resulting topology. Mis-specification of the model, or individual parameters, can lead to a well supported but erroneous result. While models are a simplistic approximation of the underlying biological drivers of evolutionary processes, getting as close an approximation as possible is essential (Box, 1979). However under- and over-parameterization have been shown to impact topology and PPs to varying degrees, in and outside the Felsenstein zone (Lemmon and Moriarty, 2004). Marginal likelihood comparison penalizes for over-parameterization and can be used to compare the fit of one model compared to another for a given data set (Xie et al., 2010). To compare how well concatenation vs. MSC fits the single copy gene data set used in this study, stepping stone comparison was conducted using the model-selection package in BEAST2 (Bouckaert et al., 2019).

### Comparison to commonly employed models and data sets

#### Phylogenetic inference using ribosomal genes

Using LSU or SSU rDNA regions for phylogenetics is common practice, at times supplemented with a small number of other genes (Shalchian-Tabrizi et al., 2006; Zhang et al., 2007; Saldarriaga et al., 2004; Murray et al., 2005; Hoppenrath and Leander, 2010). It is important to acknowledge that these represent the evolutionary history of highly conserved genes, which does not necessarily represent the species evolution and assumptions of their congruence is statistically inadequate (Degnan and Rosenberg, 2009). Yet, because rDNA sequencing is easy and inexpensive it continues to be employed for the Gonyaulacales even if it does not yield comprehensive results. Comparing the topology from a rDNA ML inference with the single gene copy MSC phylogeny presented here (Fig. 6) shows that most clades in both topologies were completely or very well supported. Within the genera *Gambierdiscus* and *Ostreopsis*, the species resolution differed between the two data sets. In several cases, the placement of sister taxa was incongruous between the two analyses. For example, the rDNA concatenation data set places *Ceratium* & *Gonyaulax* as well as *Alexandrium* and *Gambierdiscus* as sister taxa, while the single copy gene data set under MSC places *Gonyaulax* with *Protoceratium* and *Gambierdiscus* with *Pyrodinium*. This is an example of how using rDNA segments as a proxy for species evolution produces different results than an analysis of single-copy protein coding genes.

**Figure 6.**
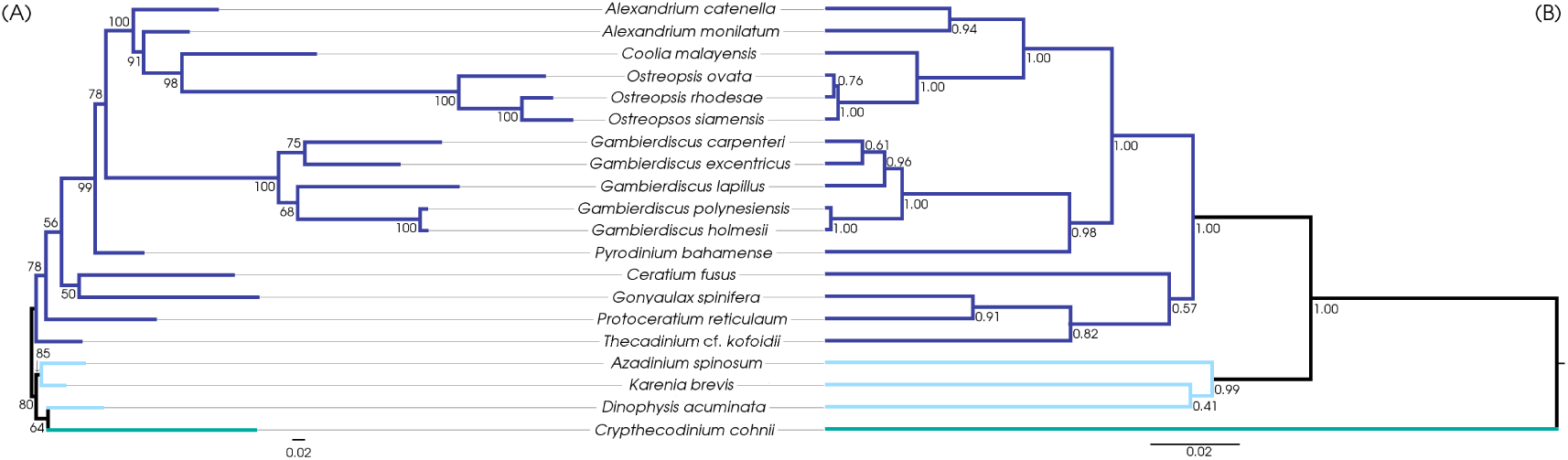
Tanglegram showing the topological differences in phylogenies from (A) concatenated rDNA genes (SSU and D1-D3 LSU) inferred with ML; and (B) MSC inference with 58 single copy genes. Gonyaulacales (#16) in purple, outgroups (#3) in light blue and taxa *incertae sedis* (#1) in teal.

#### Concatenating selected genes and using ML methods for species inference

Concatenation of alignments coupled with ML inference is a commonly used method as it is less computationally demanding than BI methods. However as demonstrated by Kubatko and Degnan (2007) and Roch and Steel (2015), this approach is error prone. Concatenation assumes uniform evolutionary history across genes, with a small amount of variation possible - however this still averages the evolutionary rate for all the input genes which doesn’t allow for divergent gene histories (Roch and Steel, 2015). The combination of concatenation and ML for phylogentic inference can result in high bootstrap values for incorrectly resolved clades, over inflating confidence in erroneous topologies (Degnan and Rosenberg, 2009). The application of concatenation in combination with ML is common practice in phylogenetic studies for gonyaulacoids (Shalchian-Tabrizi et al., 2006; Zhang et al., 2007; Saldarriaga et al., 2004; Murray et al., 2005; Hoppenrath and Leander, 2010). We investigated whether the use of a technique explicitly designed to handle multiple genes to estimate species trees would yield different results than concatenation and ML. A comparison between a BI inference under MSC and concatenated ML inference on the same single copy gene data set showed differences in topology (Fig. 7). The species resolution within the genera *Alexandrium, Gambierdiscus* and *Ostreopsis* matched between the two inference methods. The major difference was in the *Pyrodinium* placement, where the BI MSC approach places the genus sister to *Gambierdiscus* while the concatenated ML approach places it with *Gonyaulax* and *Protoceratium*. Further, the deeper branches of the phylogenies differ. The BI MSC method clusters *Ceratium, Gonyaulax, Protoceratium* and *Thecadinium* as a clade, while the concatenated ML approach clusters *Gonyaulax, Protoceratium* and *Pyrodinium* as a clade to which *Thecadinium* and then *Ceratium* feature as ancestral genera.

**Figure 7.**
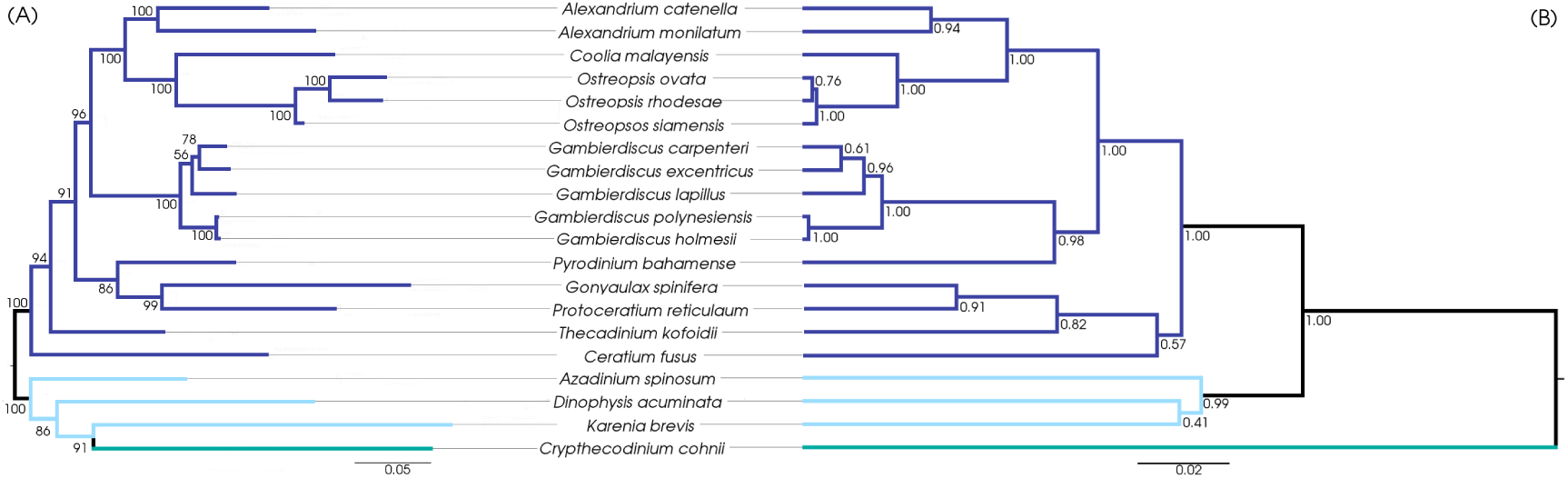
Tanglegram showing the topological differences in phylogenies with same 58 single copy gene alignments as input. (A) concatenated ML inference; and (B) MSC *BEAST2 inference. Gonyaulacales (#16) in purple, outgroups (#3) in light blue and taxa *incertae sedis* (#1) in teal.

#### Concatenating selected genes and using BI methods for species inference

Even within a BI framework concatenation can introduce a number of errors. Under simulated data sets, even under the coalescent methods, the species tree topology is inaccurate when concatenation is used (Kubatko and Degnan, 2007). Further to that, the PP values tend to be overestimated for concatenation (Suzuki et al., 2002). Theoretically for the Gonyaulacales, and taxa prone to paralogy and convoluted evolutionary histories, the MSC is a preferable approach to concatenation as MSC is more robust to ILS and LBA artifacts as well as pseudoorthologs (Liu et al., 2014; Du et al., 2019). To isolate the effects of phylogenetic model from those of the statistical framework (ML vs BI), the single copy gene data set was run with BI both under MSC and with concatenation (Fig. 8). We then used a statistical framework to compare the two model approaches to verify the veracity of model adequacy through stepping stone sampling. Stepping stone is a method for estimating marginal likelihoods of phylogenetic models, enabling model comparison and selection of the model with the better fit (Xie et al., 2011; Baele et al., 2012). The marginal likelihood of the MSC model (−160538.6) was over 10,000 log units higher than that of the concatenated single copy gene model (−170866.6), favoring the MSC approach significantly. The large difference in marginal likelihood between the models could be in part due to the inclusion of pseudoorthologs in the dataset, against which MSC models are more robust than the concatenation approach (Du et al., 2019; Roch and Steel, 2015). The resolution of *Alexandrium, Coolia* and *Ostreopsis* was identical between the two methods. Further, the species resolution within the genera *Gambierdiscus* and *Ostreopsis* was also identical 8. Differences were found in the topology, in that *Pyrodinium* clustered with *Gambierdiscus* in the MSC analysis, while for concatenation this genus clusters with *Gonyalax* and *Protoceratium* 8. The *Pyrodinium* placement also differed to the study by Price and Bhattacharya (2017) (Fig. 5), where the genus was more closely related to *Alexandrium* rather than *Gonyaulax* and *Protoceratium* in the BI topology. Further, in the MSC analysis *Ceratium, Gonyalax, Protoceratium* and *Thecadinium* formed their own clade while with concatenation, *Ceratium* and *Thecadinium* were ancestral genera to the rest of the Gonyaulacales. There was a marked difference in the internal branch arrangement, which resulted in different taxa clustering, between the concatenation and MSC methods. The concatenated approach closely mirrored the ML arrangement of taxa, apart from *Crypthecodinium* placement. Both inferences were topologically distinct to the MSC approach.

**Figure 8.**
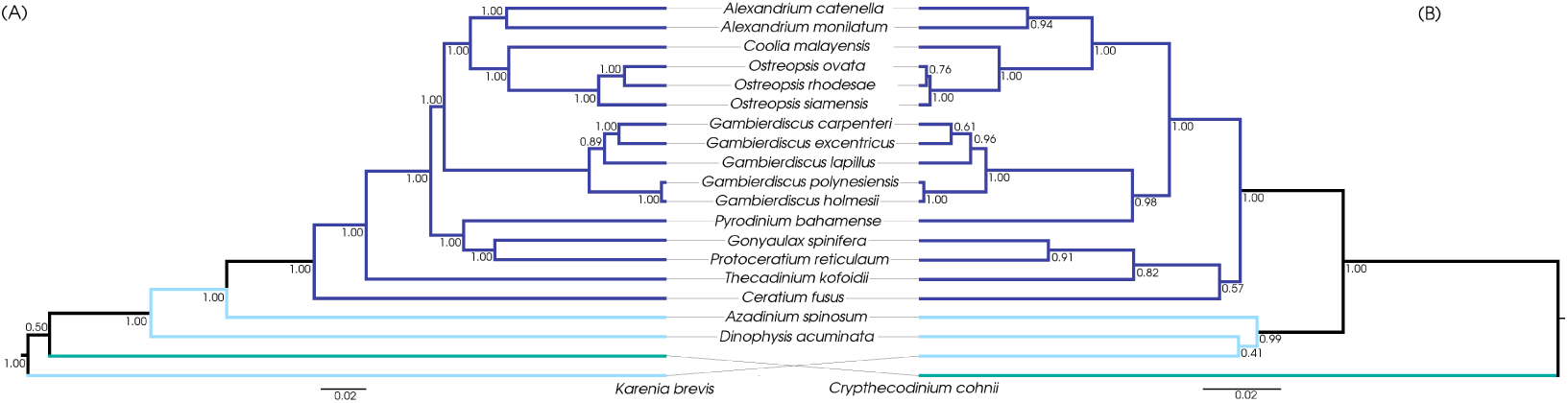
Tanglegram showing the topological differences in phylogenies with same 58 single copy gene alignments as input. (A) concatenated BEAST2; and (B) MSC *BEAST2 inference. Gonyaulacales (#16) in purple, outgroups (#3) in light blue and taxa *incertae sedis* (#1) in teal.

### Areas for possible improvement of this study

In the previous section we identified potential problems with common approaches to species inference in the literature, and in particular for the Gonyaulacales. We then sought to evaluate the effects of different methodological approaches on analytical results in the Gonyaulacales. There are several important limitations to our study.

#### Contamination of other taxa

The 650+ RNA extract submission to MMETSP was from a large number of investigators and low level contamination is inherent in the project’s data set (Keeling et al., 2014). As the cultures tested in all the studies contributing to this data set were not axenic, contamination could be bacterial or eukaryotic in nature. While any contaminating bacterial genes in our data would likely be heavily diverged and therefore obvious, eukaryotic contamination may be more subtle.

#### No representative genome for comparison

Without an available reference genome, it is difficult to evaluate the accuracy of the transcriptome assembly and whether the genes selected are single copies, or misassemblies of paralogs.

#### Different methods for RNA-seq

Three different approaches for RNA-seq library generation were employed for the libraries used in this study, the MMETSP taxa were sequenced on HiSeq platform with 50nt reads; while all other taxa were sequenced on the NextSeq platform with 75nt or 150nt reads. The different sequencing methods may each influence the single copy gene coverage and transcriptome assembly accuracy, leading to systematic error and batch effects on some taxa.

#### Total evidence phylogenetics

The method presented here purely considered the information contained in the genetic aspect of the organisms examined. Morphological characters, if evolutionarily relevant ones can be identified, and fossil dates can add another dimension to the phylogenetic inference and put the evolution within a relative time frame (Gavryushkina et al., 2017).

## CONCLUSION

This study presentse a workflow for species tree inference that implemented what is currently thought to be the best practice methods. The scripts processed RNA-seq libraries through assembly, single copy gene selection to alignment for phylogenetic species inference. As a case study exemplifying organisms rife with paralogs and ancient lineages, the Gonyaulacales were selected. The resulting phylogeny showed a well resolved, well supported inference of the Gonyaulacales evolution. This was then compared to phylogenies inferred from commonly utilized methods in the literature, and potential issues arising from these methods were discussed. By presenting a statistically rigorous method and demonstrating how it overcomes common problems in phylogenetic studies, we hope that in the future such robust, reproducible, open-access approaches to process large data-sets such as the MMETSP database can become standard practice.

## ACKNOWLEDGMENTS

The GVL section of this study was conducted inside the National eResearch Collaboration Tools and Resources (NeCTAR) research cloud, an initiative by the National Research Infrastructure for Australia (NCRIS). Gratitude to the Stanley Watson foundation, the Linnaean Society of New South Wales, and the ABRS National Taxonomy Research Student Travel Bursary for funding A. L. Kretzschmar’s attendance at the Molecular Evolution workshop at the Marine biological laboratory, Woods Hole, MA, USA. Shout out to the Taming the BEAST organizers & fellow attendees for a most illuminating workshop in February 2017 on BEAST methodology, and to Geneious for subsidizing A. L. Kretzschmar’s attendance fee. The transcriptomic sequencing was funded by A. L. Kretzschmar’s student funding and in part by an ARC Future Fellowship to S. Murray. A. L. Kretzschmar’s PhD stipend was funded through a UTS Doctoral scholarship.

## SUPPLEMENTARY MATERIAL

**Table S1:** Culturing conditions for species processed for this study.

| Species | Strain | Temp | Source location |
| --- | --- | --- | --- |
| *Gambierdiscus carpenteri* | UTSMER9A | 17 | Merimbula, AU |
| *Gambierdiscus lapillus* | HG4 | 27 | Heron Island, AU |
| *Gambierdiscus polyne-siensis* | CG15 | 27 | Rarotonga, COK |
| *Gambierdiscus holmesii* | HG5 | 27 | Heron Island, AU |
| *Thecadinium kofoidii* | THECA | 18 | Gordons bay, Sydney, AU |

**Table S2:** Transcriptomes used for study along including strain ID, source and BUSCOv2 information. MMETSP abbreviation for marine Microbial eukaryotic transcriptome sequencing project, by Moore Foundation.

| Species | Strain | complete BUSCOs | single complete BUSCOs | fragmented BUSCOs | Source |
| --- | --- | --- | --- | --- | --- |
| Gonyaulacales transcriptomes |  |  |  |  |  |
| *Alexandrium catenella* | OF101 | 110 | 74 | 3 | MMETSP0790 (Keeling et al., 2014) |
| *Alexandrium monilatum* | JR08 | 107 | 74 | 3 | MMETSP0093 (Keeling et al., 2014) |
| *Ceratium fusus* | PA161109 | 121 | 81 | 4 | MMETSP1074 (Keeling et al., 2014) |
| *Coolia malayensis* | MAB | 138 | 100 | 1 | (Verma et al., 2019) |
| *Cryptothecodinium cohnii* | Seligo | 126 | 98 | 0 | MMETSP0326.2 (Keeling et al., 2014) |
| *Gambierdiscus carpenteri* | UTSMER9A | 101 | 83 | 2 | This study |
| *Gambierdiscus excentricus* | VGO790 | 88 | 83 | 4 | (Kohli et al., 2017) |
| *Gambierdiscus lapillus* | HG4 | 141 | 98 | 2 | This study |
| *Gambierdiscus polyne-siensis* | CG15 | 104 | 81 | 3 | This study |
| *Gambierdiscus holmesii* | HG5 | 134 | 87 | 2 | This study |
| *Gonyaulax spinifera* | CCMP409 | 83 | 53 | 2 | MMETSP1439 (Keeling et al., 2014) |
| *Ostreopsis ovata* | HER27 | 132 | 99 | 2 | (Verma et al., 2019) |
| *Ostreopsis rhodesae* | HER26 | 131 | 98 | 1 | (Verma et al., 2019)) |
| *Ostreopsis siamensis* | BH1 | 132 | 98 | 1 | (Verma et al., 2019) |
| *Protoceratium reticulatum* | CCCM535=CCMP1889 | 108 | 72 | 5 | MMETSP0228 (Keeling et al., 2014) |
| *Pyrodinium bahamense* | pbaha01 | 119 | 897 | 2 | MMETSP0796 (Keeling et al., 2014) |
| *Thecadinium kofoidii* | THECA | 93 | 70 | 5 | This study |
| Outgroup transcriptomes |  |  |  |  |  |
| *Azadinium spinosum* | 3D9 | 1.8 | 81 | 4 | MMETSP1036.2 (Keeling et al., 2014) |
| *Dinophysis acuminata* | DAEP01 | 117 | 74 | 2 | MMETSP0797 (Keeling et al., 2014) |
| *Karenia brevis* | CCMP2229 | 115 | 85 | 2 | MMETSP0030 (Keeling et al., 2014) |

**Table S3:** Accession numbers for ribosomal DNA sequences used for Fig. 1. Sequences sourced from NCBI, except accesion numbers with ‘*’ sourced from the Silva database. Genes not publically available are denoted by ‘-’.

| Species | SSU seq. | D1-D3 LSU seq. |
| --- | --- | --- |
| Gonyaulacales taxa |  |  |
| *Alexandrium catenella* | AB088286 | AB088238 |
| *Alexandrium monilatum* | AY883005 | - |
| *Ceratium fusus* | AF022153 | AF260390 |
| *Coolia malayensis* | HQ897279* | KX589143 |
| *Crypthecodinium cohnii* | M64245 | - |
| *Gambierdiscus carpenteri* | EF202908 | EF202938 |
| *Gambierdiscus excen-tricus* | GETL01000157* | HQ877874 |
| *Gambierdiscus lapil-lus* | KU558930 | - |
| *Gambierdiscus poly-nesiensis* | EF202907 | This study |
| *Gambierdiscus holmesii* | This study | this study |
| *Gonyaulax spinifera* | AF022155 | DQ151558 |
| *Ostreopsis ovata* | AF244939 | KJ781420 |
| *Ostreopsis rhodesae* | KX055855 | KX055845 |
| *Ostreopsis siamensis* | KX055868 | HQ414223 |
| *Protoceratium reticu-latum* | AF274273 | EF613362 |
| *Pyrodinium ba-hamense* | AY456115 | AB936757 |
| *Thecadinium kofoidii* | AY238478 | KT371445 |
| Outgroup taxa |  |  |
| *Azadinium spinosum* | JN680857 | JN165101 |
| *Dinophysis acimu-nata* | AJ506972 | EF613351 |
| *Karenia brevis* | EF492504 | AY355458 |

